# Cell-cycle dynamics of nascent transcription and mature RNA accumulation are concordant in normal fibroblasts

**DOI:** 10.1101/2025.09.12.675830

**Authors:** Melanie Gucwa, Kohta Ikegami

## Abstract

Cell-cycle dynamics of gene expression are fundamental to life, yet their origin remains unclear. A prevailing model, derived from cancer cells, posits that transcription occurs in one cell cycle phase, while mature RNA accumulates in subsequent phases, suggesting temporal segregation of transcriptional regulation and RNA abundance. Whether this paradigm applies to normal human cells is unclear. Here, we co-profiled nascent transcription and mature RNAs in cycling human fibroblasts. The two dynamics were strongly concordant, with no evidence of a lag extending cell-cycle phases. Our data suggest a widespread transcription-to-maturation lag is not a general feature of human cells.

## BACKGROUND

Precise temporal control of gene expression during the cell cycle is essential for cellular homeostasis. Disruption of this regulation can drive uncontrolled proliferation and tumorigenesis [1]. More than 1,000 human genes show cell cycle phase-specific accumulation of transcripts [2–7]. However, the transcriptional mechanisms that generate these dynamics remain controversial. One prevailing model proposes a transcription-to-maturation lag, in which transcription occurs in one cell-cycle phase, but mature transcripts accumulate in the subsequent phases [8]. For example, transcripts produced in early G1 accumulate in S phase, those produced in G1/S accumulate at M phase, and those produced in M phase accumulate in G0/G1 [8]. This model, based on MCF7 cancer cells arrested at defined cell cycle phases, implies that transcriptional regulation and transcript abundance are temporally segregated during the cell cycle. An alternative model, supported by studied of classic cell cycle-regulated genes such as E2F targets, suggests that transcription and transcript accumulation occur within the same cell-cycle phase [9]. Because few studies have measured nascent and mature transcripts simultaneously, the temporal relationship between transcription and transcript accumulation remains unresolved. A recent single-cell study in HEK293T cells reported only a subtle lag along a pseudo-time scale, that did not extend across cell-cycle phases, likely reflecting transcriptional elongation time [3]. Whether a pervasive transcription-to-maturation lag exists in non-transformed cells remains unknown.

## RESULTS AND DISCUSSION

To investigate cell-cycle dynamics of transcription and transcript accumulation in cycling non-transformed cells, we introduced the Fucci4 cell-cycle fluorescent reporters [10] into the hTERT-immortalized human fibroblast cell line BJ-5ta (BJ-Fucci) (**Fig. 1A**). The Fucci4 system consists of four components: Cdt1^aa30-120^ fused to mKusabira-Orange2 (mKO2) which accumulates in G1; SLBP^aa18-126^ fused to mTurquoise2, which accumulates in G1 and S; Geminin^aa1-110^ fused to Clover, which accumulates in S and G2; and histone H1 fused to mMaroon1, which visualizes chromatin [10]. We validated BJ-Fucci cells by time-lapse fluorescent imaging (**Fig. S1A; Table S1; Movie 1**). As expected, cells lacked mKO2, mTurquoise2, and Clover during early G1, showed peak mKO2 expression during G1, co-expressed all three reporters in early S, and showed peak Clover expression during late-S/G2/M (**Fig. 1B, S1B**).

**Figure 1.**
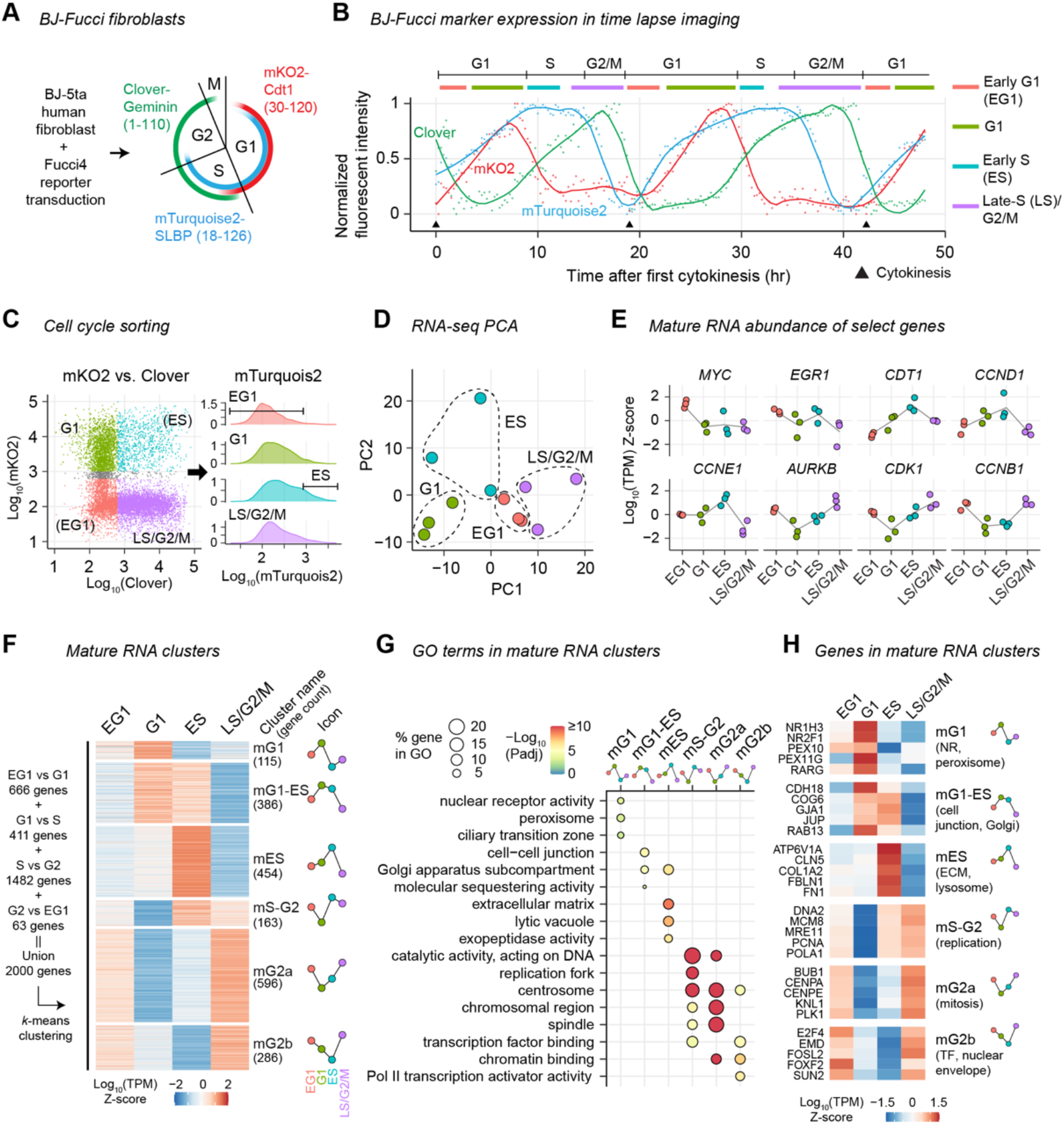
Cell-cycle dynamics of mature RNA in BJ-Fucci fibroblasts. **(A)** BJ-Fucci fibroblasts. BJ-5ta fibroblast cells are transduced with the Fucci4 cell-cycle reporters mKO2-Cdt1^aa30-120^, mTurquoise2-SLBP^aa18-126^, Clover-Geminin^aa1-110^, and mMaroon1-H1. **(B)** Nuclear fluorescence signals in a single BJ-Fucci cell (0–19 hr), daughter cell (19–43 hr), and grand-daughter cell (43hr–) measured by time-lapse imaging (duration 48 hr, interval 15 min). Circle, min-max normalized log_10_ intensity. Line, Loess regression. Top, empirical cell cycle phase annotations. See **Figure S1** for additional cells. **(C)** FACS of BJ-Fucci cells into early G1 (EG1), G1, early S (ES), and late S (LS)/G2/M phases. Populations indicated in the Clover/mKO2 plot (left) are shown in mTurquoise2 histogram (right). EG1 and ES are subsets of Clover/mKO2 populations defined by mTurquoise2 intensity. **(D)** Principal component analysis (PCA) of RNA-seq biological replicates by their mature RNA abundance (log_10_ RNA-seq TPM) in all genes. **(E)** Mature RNA abundance (gene-level z-score of log_10_ RNA-seq TPM) of select genes known to be dynamic in expression during the cell cycle. **(F)** K-means clustering of the union of differentially expressed genes by their mature RNA dynamics (gene-level z-score of log_10_ RNA-seq TPM). “Icons” are within-cluster mean of log_10_ TPM z-score and used in subsequent figures to graphically represent cluster dynamics. **(G)** Molecular Function and Cellular Component GO terms enriched in dynamic mature RNA clusters. Top 3 most enriched terms from each cluster are selected and their enrichment in each cluster was shown. **(H)** Mature RNA abundance dynamics (log_10_ RNA-seq TPM z-score) of five representative genes in each cluster.

To determine the dynamics of mature RNA abundance during the cell cycle, we sorted BJ-Fucci cells into four phases using fluorescence-activated cell sorting (FACS): early G1 (mKO2^−^Clover^−^ mTurquoise2^−^), G1 (mKO2^+^Clover^−^), early-S (mKO2^+^Clover^+^mTurquois^+^), and late-S/G2/M (mKO2^−^ Clover^+^) (**Fig. 1C; Fig. S2A; Table S2**). We then sequenced poly(A)-selected RNAs using RNA-seq from each population (**Fig. S3A**). We confirmed high concordance between biological replicates, as well as expected differences between cell cycle phases (**Fig. 1D; Fig. S3B**). The RNA-seq data recapitulated known cell-cycle dynamics of mature RNA abundance, including early-G1 expression of *MYC* [11] and *EGR1* [12], G1-to-early-S elevation of *CDT1* and *CCND1* [13,14], peak early-S expression of *CCNE1* [13], and late-S/G2/M expression of *AURKB, CCNB1*, and *CDK1* [13,15] (**Fig. 1E**). Next, we identified genes whose RNA abundance differed significantly between neighboring cell-cycle phases. There were 666 differentially abundant genes between early G1 and G1, 411 between G1 and early S, 1,482 between early S and late S/G2/M, and 63 between late S/G2/M and early G1, totaling 2,000 unique genes (**Fig. S3C**). We then clustered these 2,000 genes based on RNA abundance dynamics across the cell cycle and identified six distinct patterns (**Fig. 1F; Table S3**). The “mG1” cluster (*m* for mature RNA) had peak abundance in G1 (115 genes). The “mG1-early S (ES)” cluster had peak abundance in both G1 and early-S (386 genes), while the “mES” cluster had one peak in early-S (454 genes). The “mS-G2” cluster had peak abundance in early-S and late S/G2/M (163 genes). Finally, the “mG2a” (596 genes) and “mG2b” (286 genes) clusters had a primary peak in late-S/G2/M and a secondary peak in early G1 (**Fig. 1F**). Gene ontology analysis indicated that these clusters were enriched for functionally distinct classes of genes (**Fig. 1G**). The mG1 cluster was enriched for nuclear receptors (e.g. *NR2F1, RARG*) and peroxisome components (e.g. *PEX10, PEX11G*) (**Fig. 1G, H**). The mG1-ES and mES clusters were enriched for fibroblast-related functions, such as cell-cell junctions and adhesion (e.g. *GJA1, JUP*; mG1-ES) and extracellular matrix components (e.g. *COL1A2, FN1*; mES). The mS-G2 cluster was strongly enriched for DNA replication-related genes (e.g. *PCNA, MRE11*), while mG2a was enriched for mitosis-related genes (e.g, *PLK1, CENPA*), and mG2b cluster was enriched for transcription factors and nuclear envelope factors (e.g. *E2F1, SUN2*). Notably, no cluster peaked in early G1. Together, these results defined the cell cycle dynamics of mature RNA abundance in normal fibroblasts, setting the stage for investigating their relationship to transcription timing.

To define nascent transcriptional dynamics, we sorted BJ-Fucci cells in early G1, G1, and late S/G2/M phases and performed GRO-seq (early-S phase was excluded due to insufficient cells) (**Fig. S4A**). GRO-seq quantifies nascent transcripts produced during a short incubation of permeabilized cells with bromouridine (BrU) by sequencing BrU-incorporated RNAs [16]. We confirmed strand-specific GRO-seq signals originating from both exons and introns, a feature of nascent transcripts (**Fig. 2A**). The GRO-seq signals were highly concordant between biological replicates, yet distinct across cell cycle phases (**Fig. 2B; Fig. S4B**). We identified genes whose nascent transcription levels (GRO-seq read counts) differed significantly between neighboring cell-cycle phases. There were 583 differentially transcribed genes between early G1 and G1, 2,622 between G1 and late-S/G2/M, and 475 between late S/G2/M and early G1, totaling 2,771 unique genes (**Fig. S4C**). Clustering these genes by nascent transcription dynamics identified six clusters (**Fig. 2C; Table S4**). The “tEG1” cluster (*t* for transcription) showed peak transcription in early G1, followed by a stepwise decline in G1 and late S/G2/M. This cluster was enriched for components of the Cdc45-MCM-GINS (CMG) complex involved in replication origin licensing (e.g. *MCM2, MCM3, GINS2*), a process that occurs during G1 prior to S phase [17] (**Fig. 2D, E**). The “tG1a”, “tG1b” and “tG1c” clusters exhibited peak transcription at G1 and were enriched for fibroblast-related functions, such as cell projection (e.g. *BIN1*; tG1a), adhesion (e.g. *CDH2*; tG2b), and extracellular matrix (e.g. *COL1A1*; tG1c). The “tG2a” and “tG2b” clusters had peak transcription in late S/G2/M and were strongly enriched for DNA replication (e.g. *DNMT1*; tG2b) or mitosis-related genes (e.g. *CCNB1*; tG2a and tG2b).

**Figure 2.**
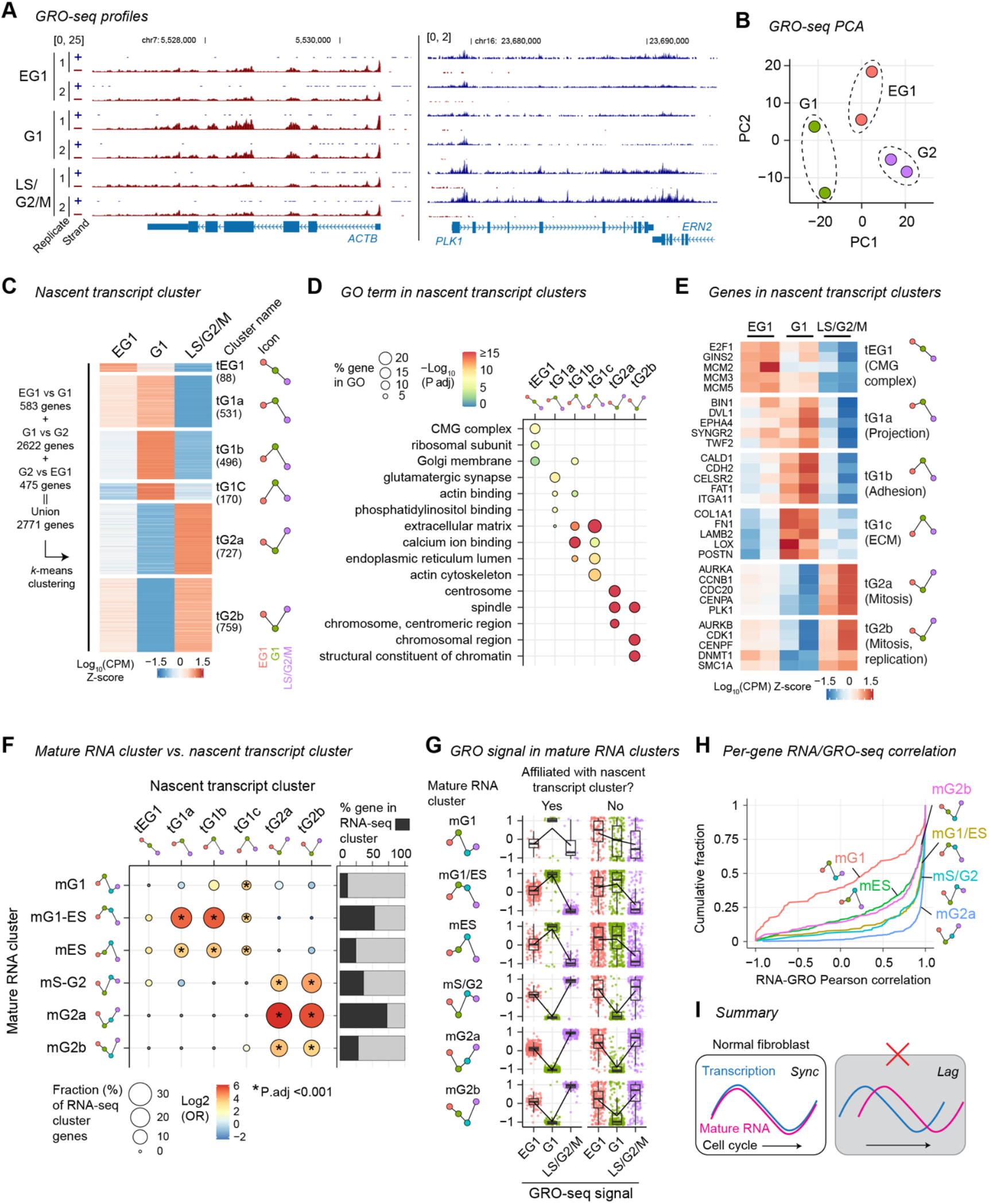
Strong concordance between the cell-cycle dynamics of mature RNA abundance and nascent transcription. **(A)** GRO-seq read density, normalized to sequencing depth, at the *ACTB* (left) and *PLK1* (right) loci. Signals for the plus and minus strands are separately shown. Numbers in brackets indicate Y-axis signal range. **(B)** PCA of GRO-seq biological replicates by their nascent transcript levels (log_10_ GRO-seq CPM) in all genes. **(C)** K-means clustering of the union of differentially transcribed genes by their nascent transcript dynamics (gene-level z-score of log_10_ GRO-seq CPM). “Icons” are within-cluster mean of log_10_ CPM z-score and used in subsequent figures to graphically represent cluster dynamics. **(D)** Molecular Function and Cellular Component GO terms enriched in dynamic nascent transcript clusters. Top 3 most enriched terms from each cluster are selected and their enrichment in each cluster was shown. **(E)** Nascent transcript dynamics (log_10_ GRO-seq CPM z-score) of five representative genes in each cluster. **(F)** Gene overlap between mature RNA clusters and nascent transcript clusters. Left, Fraction of genes in mature RNA clusters affiliated with nascent transcript clusters. Circle color indicates odds ratio (OR) of overlap, with Fisher’s exact test P-value indicated by asterisks. Right, Total fraction of genes in mature RNA clusters affiliated with any nascent transcript cluster. **(G)** Nascent transcript dynamics (log_10_ GRO-seq CPM z-score) of genes stratified by mature RNA clusters (rows) and whether they are affiliated with any one of nascent transcript clusters (column). Overlaid boxplots indicate interquartile (IQR) range with whiskers extending to farthest data points within 1.5x IQR. Lines connect means. **(H)** Cumulative distribution of per-gene Pearson correlation coefficients between mature RNA abundance and nascent transcription within each mature RNA cluster. Pearson correlation is computed between log_10_ RNA-seq TPM z-score and log_10_ GRO-seq CPM z-score at EG1, G1, and LS/G2/M, for each gene. **(I)** Summary. In BJ-Fucci fibroblasts, the dynamics of nascent transcription and mature RNA dynamics are strongly concordant (left) and do not exhibit a widespread lag (right).

To investigate the relationship between transcription and mature RNA abundance during the cell cycle, we assessed the overlap of gene membership between mature mRNA clusters and nascent transcription clusters. We observed a strong concordance between the two. Genes in mature RNA clusters peaking in G1 or early S (mG1, mG1-ES, and mES) were overrepresented in transcriptional clusters with peaks in G1 (tG1a, tG1b, tG1c) (**Fig. 2F**). Likewise, significant number of genes overlapped between mature RNA clusters and transcriptional clusters with peaks in late S/G2/M. The strongest overlap was observed in late-S/G2/M, where 72% of mG2a genes belonged to either tG2a or tG2b (**Fig. 2F**). An exception was the mG1 mature RNA cluster, in which most genes did not exhibit dynamic nascent transcription, suggesting a larger contribution of post-transcriptional regulation to mG1 mature RNA dynamics. Importantly, we found no significant overlap between G1/early S-peaked mature RNA clusters and late S/G2/M-peaked transcriptional clusters, or vice versa. These results indicate that transcription dynamics and mature RNA dynamics are aligned within cell-cycle phases, with no evidence of a systematic transcription-to-maturation lag extending across cell-cycle phases.

We observed concordance beyond the specific genes shared between mature RNA clusters and transcriptional clusters. Genes in the dynamic mature RNA clusters exhibited concordant nascent transcription patterns regardless of their assignment to specific transcriptional clusters, except those in the mG1 cluster (**Fig. 2G**). We further computed correlations between mature RNA and nascent transcript levels across the cell cycle for each gene. Approximately 75% of mature RNA cluster genes, excluding mG1 cluster genes, showed a strong positive correlation (Pearson’s *r* > 0.5) (**Fig. 2H**). Taken together, our data indicated that mature RNAs enriched in G1 or G2 phases were transcribed in the same G1 or G2 phase, and not in a preceding phase.

## CONCLUSIONS

Our analysis revealed a strong consistency between the cell cycle dynamics of nascent transcription and that of mature RNA abundance in normal fibroblasts. We did not find evidence for a widespread transcription-to-maturation lag extending cell-cycle phases, similar to a recent report in HEK293T cells [3]. This conclusion differs from an observation of a pervasive lag in a cancer cell line MCF7 [8]. One potential reason for this difference is that such a lag may be a form of dysregulation in certain cancer cells, contributing to rapid proliferation. Another reason may be a technical difference: while the previous study investigated cells arrested at specific cell-cycle phases, this present study analyzed unperturbed, naturally cycling cells. Together, these findings indicate that a widespread transcription-to-maturation lag is not a general feature of human cells.

**Figure S1.**
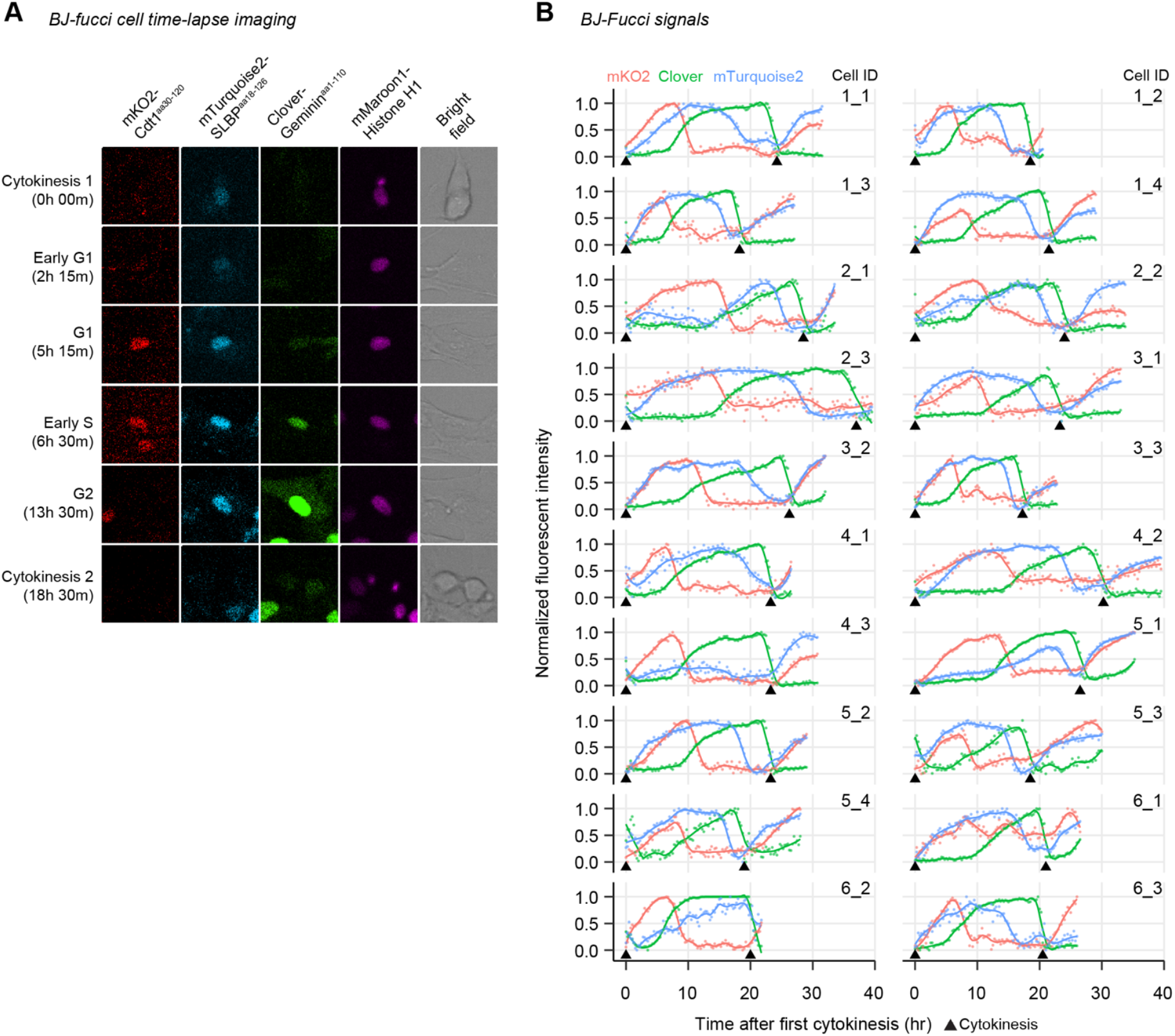
Time-lapse imaging of BJ-Fucci cells. **(A)** A representative BJ-Fucci cell in time-lapse confocal imaging (duration 48 hr, interval 15 min). Fluorescent signal of mKO2, mTurquoise2, Clover, and mMaroon1 and bright-field image for a single fibroblast at indicated cell cycle stage. **(B)** Nuclear fluorescence signals in individual BJ-Fucci cells. Each plot shows one cell cycle and a part of the following cycle of the daughter cell. Circle, min-max normalized log_10_ intensity. Line represents Loess regression. Arrowhead indicates cytokinesis of the daughter cells.

**Figure S2.**
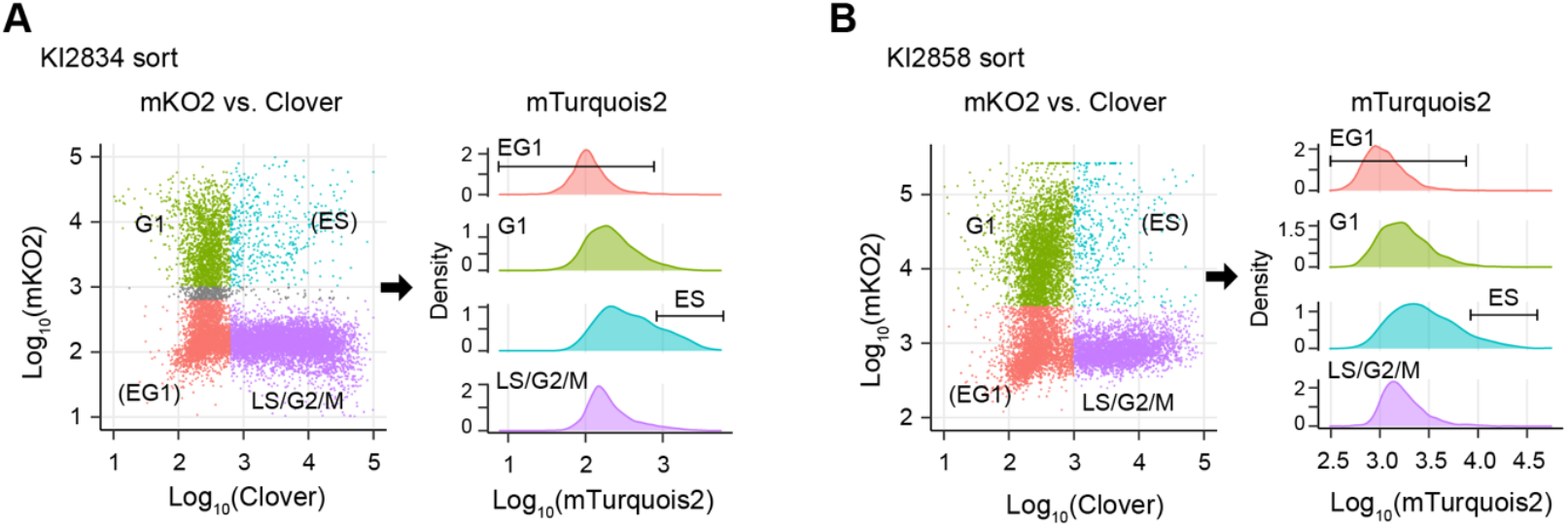
FACS of BJ-Fucci cells. **(A, B)** FACS of BJ-Fucci cells into early G1 (EG1), G1, early S (ES), and late S (LS)/G2/M phases. Two biological replicates (A and B) of FACS experiments are shown. Another replicate is shown in Figure 1. Populations indicated in the Clover/mKO2 plot (left) are shown in mTurquoise2 histogram (right). EG1 and ES are subsets of Clover/mKO2 populations defined by mTurquoise2 intensity.

**Figure S3.**
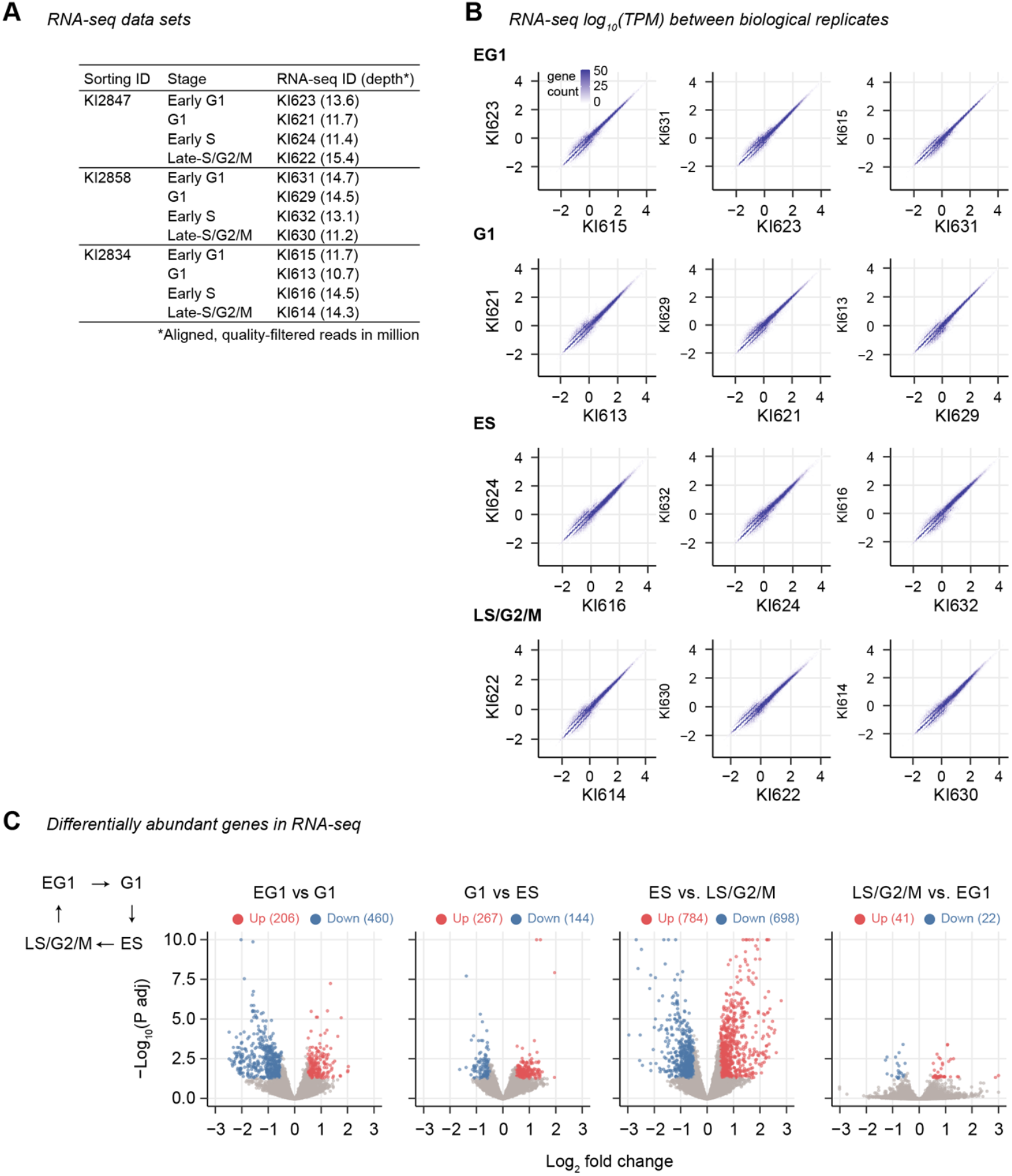
RNA-seq analysis of cell cycle-sorted BJ-Fucci cells. **(A)** RNA-seq data sets produced in this study. **(B)** Pair-wise comparison of RNA-seq signals (log_10_TPM) in each cell-cycle phase between biological replicates. **(C)** Differential expression analysis between neighboring cell-cycle phases. Left diagram shows the cell cycle phase order, indicating the sequential phases compared in each plot. Volcano plots show log_2_ fold change and DESeq2 adjusted P-value representing the significance of RNA abundance difference.

**Figure S4.**
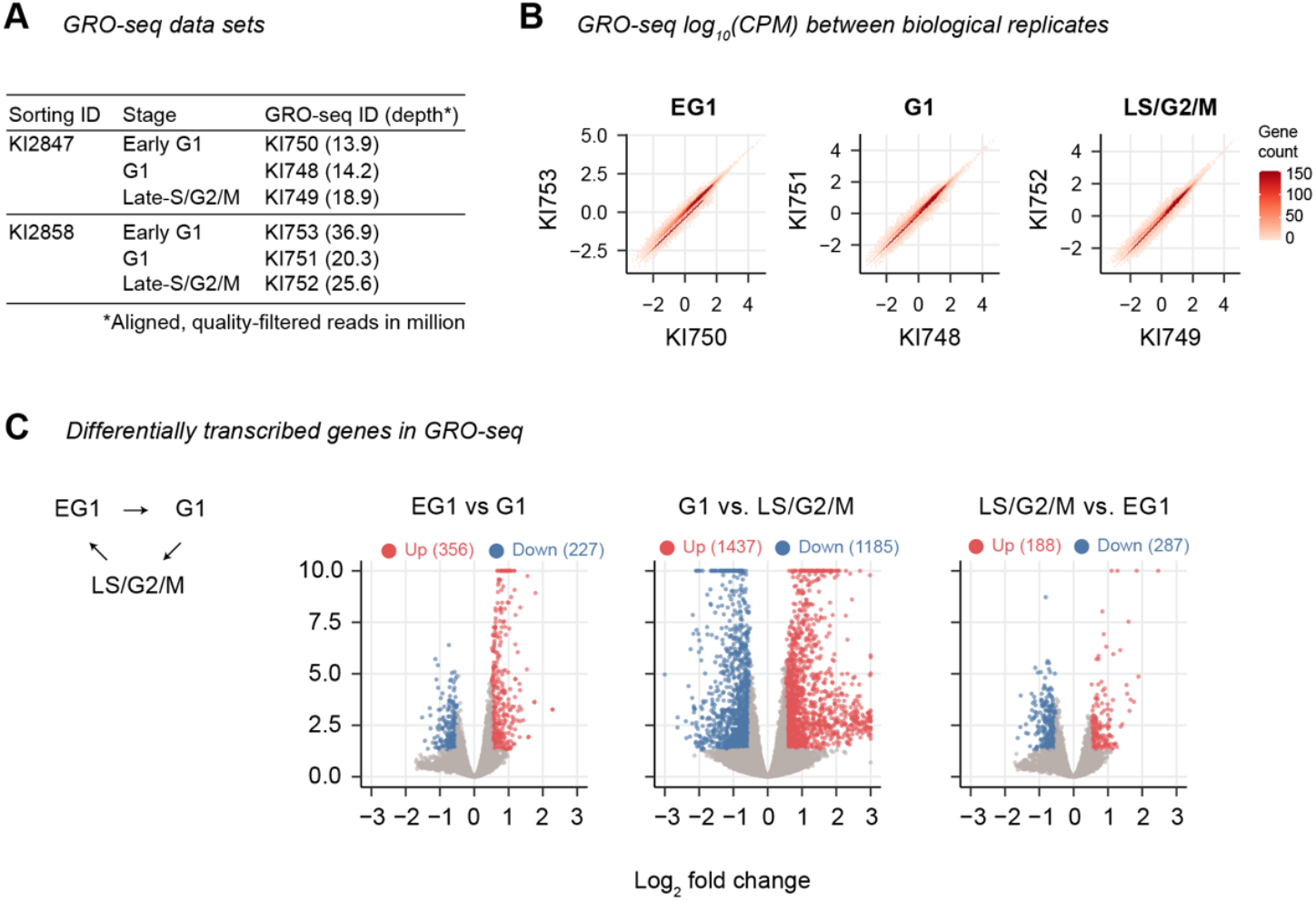
GRO-seq analysis of cell cycle-sorted BJ-Fucci cells. **(A)** GRO-seq data sets produced in this study. **(B)** Pair-wise comparison of GRO-seq signals (log_10_CPM) in each cell-cycle phase between biological replicates. **(C)** Differential expression analysis between neighboring cell-cycle phases. Left diagram shows the cell cycle phase order, indicating the sequential phases compared in each plot. Volcano plots show log_2_ fold change and DESeq2 adjusted P-value representing the significance of nascent transcription difference.

## METHODS

### Establishment of BJ-Fucci cell line

The hTERT-immortalized human dermal fibroblast cell line BJ-5ta (source: foreskin of male neonate) was purchased from ATCC (ATCC catalog # CRL-4001). BJ-5ta retained normal fibroblast growth phenotypes and did not exhibit transformed phenotypes, as described in the original publication [18]. To introduce Fucci4 fluorescent markers, we produced lentiviruses carrying Clover-Geminin^aa1-110^-IRES-mKO2-Cdt1^aa30-120^ (gift from Michael Lin; Addgene ID83841; RRID:Addgene_83841) or mTurquoise2-SLBP^aa18-126^-IRES-H1-mMaroon1 (gift from Michael Lin; Addgene ID83842; RRID:Addgene_83842), individually. A detail procedure for lentivirus production using HEK293FT cells is described in our previous publication [19]. BJ-5ta cells were co-transduced with the two lentivirus supernatants at 1:60 dilution in the presence of 6.7 µg/mL polybrene. The transduced cells were selected for their simultaneous expression of Clover and mMaroon1 and then for their simultaneous expression of mKO2 and mMaroon1 using FACS. The resultant double-selected cells are “BJ-Fucci” cells (cell ID cc1357-2ss). Independently, we transduced BJ-5ta cells with the two vectors individually (cell ID cc1376-1 for mKO2/Clover and cc1376-2 for mTurquois2/mMaroon1) or with mKO2-SLBP^aa18-126^ (gift from Michael Lin; Addgene ID 83914; RRID:Addgene_83914; cell ID cc1376-3) or with Clover-Geminin^aa1-110^ (gift from Michael Lin; Addgene ID 83915; RRID:Addgene_83915; cell ID cc1376-4), to generate references for signal compensation in flow cytometry. BJ-5ta and all its derivatives, including BJ-Fucci cells, were cultured in high-glucose DMEM (Gibco, 11965-092) containing 9% fetal bovine serum (FBS), 90 U/mL penicillin, 90 µg/mL streptomycin streptomycin at 37ºC under 5% CO_2_.

### Time lapse confocal microscopy

BJ-Fucci cells in a glass-bottom dish (MatTek P35G-1.5-20-C) were placed in a microscope stage-top incubator set to 37ºC and 5% CO_2_ (Tokai Hit) and imaged 6 areas in the dish every 15 minute for 48 hours using a Nikon AXR inverted confocal laser scanning microscope with a 20x objective. For each frame for each image area, mTurquioise2, Clover, mKO2, mMaroon1, and bright field images were obtained. The following excitation (ex) lasers and emission (em) tunable filter settings were used: mTurquioise2, ex 445 nm, em 464–497 nm; Clover, ex 488 nm, em 505–547 nm; mKO2, ex 561 nm, em 571–591 nm; mMaroon1, ex 594 nm, em 609–712 nm (**Movie 1**). To plot the cell-cycle dynamics of fluorescent signal intensities per cell, we first identified cells that exhibited two cytokinesis events in the imaging area within the imaging duration and obtained mean nuclear fluorescent intensities at each frame (**Table S1**). We then normalized the log_10_-scaled intensities for each cell such that the minimum intensity was set 0 and the maximum intensity was set 1 during the cell cycle, to use in plots.

### Fluorescence-Activated Cell Sorting (FACS)

Before sorting, fluorescence compensation was set up using BJ-5ta cell populations expressing mKO2-SLBP alone, Clover-Geminin alone, Clover-Geminin-IRES-mKO2-Cdt1 alone, or mTurquoise2-SLBP-IRES-H1-mMaroon1 alone. Approximately 20 hours before sorting, BJ-Fucci cells that reached confluency were passaged to 8–20 15-cm culture dishes at a density of 2.5 million cells per dish. Cells were dissociated from the dishes with TrypLE (Thermo Fisher), filtered through a 40-µm cell strainer, resuspended in the complete culture medium, and sorted into the culture medium using BD FACSAriaII (sorting experiment ID: KI2834 and KI2847) or BD FACSAriaIIu (KI2858) sorters. The following populations were sorted: mKO2^−^Clover^−^mTurquoise2^−^ (early G1), mKO2^+^Clover^−^ (G1), mKO2^+^Clover^+^mTurquois^+^ (early S), and mKO2^−^Clover^+^ (late-S/G2/M). The following excitation (ex) lasers and emission (em) filter settings were used: mTurquioise2, ex 405 nm, em 450/50 nm; Clover, ex 488 nm, em 530/30 nm; mKO2, ex 561 nm, em 582/15 nm; mMaroon1, ex 561 nm, em 670/30 nm (FACSAriaII) or em 670/14 (FACSAriaIIu). After sorting, cells were stored with Trizol LS (Thermo Fisher) at –80ºC for RNA-seq or directly proceeded to nuclear isolation described in the GRO-seq section. Fluorescent signals of cells immediately prior to sorting are listed in **Table S2**.

### RNA-seq

Total RNAs were purified by Trizol LS (Invitrogen 10296028) and treated with DNase I (Invitrogen Turbo DNase AM2238). mRNAs were isolated and fragmented using NEBNext Poly(A) mRNA Magnetic Isolation Module (New England Biolabs E7490). cDNAs were synthesized using the cDNA synthesis module in NEBNext UltraII Directional RNA Library Prep Kit for Illumina (NEB E7660). Sequencing libraries were prepared using NEBNext Ultra DNA Library Prep Kit (New England Biolabs, E7370). Libraries were processed on the Illumina HiSeq 2500 for single-end 50-nt sequencing.

### GRO-seq

Nuclei were isolated by incubating cells in hypotonic NP40 lysis buffer (10 mM NaCl, 3 mM MgCl2, 0.5% MP-40, 10 mM Tris, pH 7.5) supplemented with RNase Inhibitor on ice and resuspended in Nuclear Storage buffer (50 mM Tris pH 8.0, 0.1 mM EDTA, 5 mM MgCl2, 40% glycerol, RNase inhibitor). The nuclear suspension was mixed with an equal volume of 2x NRO buffer (10 mM Tris pH 8.0, 5 mM MgCl2, 1 mM DTT, 300 mM KCl, 0.5 mM ATP, 0.5 mM GTP, 0.5 mM BrUTP, 2 µM CTP). The sample was incubated without sarkosyl for 4 min at 30ºC and then with 0.5% sarkosyl for 4 min at 30ºC (total 8 min). RNAs were purified from the reaction by Trizol LS (Invitrogen 10296028) followed by isopropanol precipitation. RNAs were treated with TurboDNase (Ambion AM18907) and fragmented by Fragmentation Buffer (Ambion AM8740). BrU-incorporated RNA fragments were immunoprecipitated with mouse monoclonal anti-BrdU antibody 3D4 (BD Biosciences 555627 Lot 7033666) and used to construct DNA sequencing libraries using NEBNext Ultra II Directional RNA Library Prep kit (New England Biolabs E7760). DNA libraries were processed on an Illumina NextSeq machine for paired-end 42-nt sequencing.

### RNA-seq analysis

RNA-seq reads were aligned to the GRCh38 human reference genome with the Gencode v38 basic gene annotation (60,649 genes) [20] using STAR 2.7.9 [21] with the default alignment parameters except using “clip3pAdapterSeq AGATCGGAAGAGCACACGTCTGAACTCCAGTCA”. Because a principal component analysis (PCA) found a batch effect of cell sorting in the first principal component, raw read counts were processed in Combat-seq [22] in R to correct for the sorting batch. The corrected read counts were used to compute transcripts per million (TPM), such that TPM =10^6^ x RPK/sum(RPK), where RPK is [1+corrected read count]/[transcript length in kb], and sum(RPK) is sum of RPKs for all genes. TPMs were used to compute z-scores of log_10_(mean(TPM)) for each gene across cell-cycle stages, where mean(TPM) is the mean of TPMs in biological replicates. In principal component analysis (PCA), genes with zero read count in any one replicate were removed, and then sample-wide mean-centered log_10_TPM were used as an input in the *prcomp* function in R. Corrected read counts were used in DESeq2 program [23] in R to identify genes with different mRNA abundance between early G1 and G1, between G1 and early S, between early S and late S/G2/M, and between late S/G2/M and early G1. Genes with adjusted P-value < 0.05 and absolute log_2_ fold change > 0.5 were considered differentially expressed (**Table S3**). To identify mature RNA clusters, we applied k-means clustering (k=6) to the z-scores of log_10_(mean(TPM)) for the union of the differentially expressed genes (2,000 genes). For Gene Ontology (GO) enrichment analysis, a subset of genes in mature RNA clusters that were “protein_coding” in the gene_type annotation were used in Metascape [24] with the default setting except selecting GO Molecular Functions and GO Cellular Components as the search data base.

### GRO-seq analysis

GRO-seq read pairs with maximum fragment length of 2000 bp were aligned to the GRCh38 human reference genome using Bowtie2 with the parameter set of -X 2000 --no-mixed --no-discordant. Reads with MAPQ score greater than 20 were used in downstream analyses. The 5’-end of the paired-end fragment reflects the initial location of RNA Polymerase II that produces nascent transcripts. For each transcript in the Gencode v38 basic gene annotation (111,832 transcripts), we counted the number of the overlapping paired-end fragments at their 5’-end (1 base). To generate the genomic density profile for visualization, the fragment counting was performed at every base in the genome after extending +/–10 bp from the 5’ end. Out of the 111,832 transcripts, we selected a single transcript with the longest transcript size (sum of exons) to represent one of the 60,649 genes. The per-gene GRO-seq fragment counts were processed in Combat-seq [22] in R to correct for the sorting batch. The corrected fragment counts were used to compute counts per million (CPM), such that CPM =10^6^ x CPK/sum(CPK), where CPK is [1+corrected fragment count]/[gene length in kb], and sum(CPK) is sum of CPKs for all genes. CPMs were used to compute z-scores of log_10_(mean(CPM)) for each gene across cell-cycle stages, where mean(CPM) is the mean of CPMs in biological replicates. In PCA, genes with zero fragment count in any one replicate were removed, and then sample-wide mean-centered log_10_CPM were used as an input in the *prcomp* function in R. Corrected fragment counts were used in DESeq2 program [23] in R to identify genes with different nascent transcript abundance between early G1 and G1, between G1 and late S/G2/M, and between late S/G2/M and early G1. Genes with adjusted P-value < 0.05 and absolute log_2_ fold change > 0.5 were considered differentially transcribed (**Table S4**). To identify nascent transcript clusters, we applied k-means clustering with k=6 to the z-scores of log_10_(mean(CPM)) for the union of the differentially transcribed genes (2,771 genes). For GO enrichment analysis, a subset of genes in nascent transcript clusters that were “protein_coding” in the gene_type annotation were used in Metascape [24] with the default setting except selecting GO Molecular Functions and GO Cellular Components as the search data base.

### Joint analysis of RNA-seq and GRO-seq

The statistical significance of the gene membership overlap between mature RNA clusters and nascent transcript clusters was computed by Fisher’s exact test. The P-values were adjusted for multiple testing using the Benjamini-Hochberg Procedure. To compute correlation between RNA-seq signals and GRO-seq signals for each gene, we first removed the early S phase data point from the RNA-seq data (because there was no early-S phase data point in GRO-seq) and then recomputed the z-scores of log_10_(mean(TPM)) for RNA-seq. We then computed Pearson correlation coefficient between RNA-seq z-scores and GRO-seq z-scores at early G1, G1, and G2.

## DECLARATIONS

### Ethics approval and consent to participate

Not applicable.

### Consent for publication

All authors agreed for publication of this manuscript.

### Availability of data and materials

All genomics data are available in Gene Expression Omnibus under accession ID GSE305617. The BJ-Fucci cell line is available upon request to corresponding author under material transfer agreement.

### Authors’ contributions

Conceptualization, K.I.; methodology, M.G. and K.I.; formal analysis, K.I.; investigation, K.I.; writing – original draft, K.I.; writing – review & editing, M.G., K.I.; visualization, K.I.; supervision, K.I.; project administration, K.I.; funding acquisition, K.I.

## Acknowledgements

We thank Sebastian Pott for critical feedback on this study. We thank the genomics core facility, the flow cytometry facility, the microscopy facility at the University of Chicago and the microscopy facility at Cincinnati Children’s Hospital Medical Center for their assistance.

## SUPPLEMENTAL INFORMATION

Table S1: Fucci4 marker signals in time-lapse imaging

Table S2: Fucci4 marker signals in FACS

Table S3: RNA-seq TPM, z-score, differentially abundant genes, and clusters Table S4: GRO-seq CPM, z-score, differentially transcribed genes, and clusters

Movie 1: Time-lapse imaging of BJ-Fucci cells

